# FEDRANN: effective long-read overlap detection based on dimensionality reduction and approximate nearest neighbors

**DOI:** 10.1101/2025.05.30.656979

**Authors:** Jia-Yuan Zhang, Changjiu Miao, Teng Qiu, Xiaoshuang Xia, Lei He, Junyi He, Chunlei Yang, Yuhui Sun, Tao Zeng, Yuxiang Li, Xun Xu, Yijun Ruan, Yuliang Dong

## Abstract

Overlap detection is a key step in *de novo* genome assembly pipelines based on the Overlap-Layout-Consensus (OLC) paradigm. However, existing methods for overlap detection either rely on heuristic seed-and-extension strategies or locality-sensitive hashing (LSH), both of which struggle to handle repetitive genomic regions and the computational burden of large-scale datasets. Here, we present FEDRANN, a novel strategy for overlap graph construction that integrates feature extraction, dimensionality reduction (DR), and approximate nearest neighbor (ANN) search. We find the pipeline combining inverse document frequency (IDF) transformation, sparse random projection (SRP), and NNDescent enables accurate detection of overlaps across diverse datasets. We developed an efficient open-source implementation of this pipeline named Fedrann (https://github.com/jzhang-dev/fedrann). Through systematic benchmarking on real long-read sequencing data, we demonstrate that Fedrann produces overlap graphs comparable to or better than those generated by existing state-of-the-art tools, including MECAT2, minimap2, and wtdbg2, while maintaining competitive runtime. Despite being implemented primarily in Python, *Fedrann* achieves performance on par with tools written in compiled languages, owing to matrix-based representations and C-accelerated numerical libraries. Our results suggest that DR and ANN techniques offer a promising new direction for scalable and accurate overlap detection in long-read assembly and broader sequence similarity search tasks.

## Introduction

*De novo* assembly reconstructs a genome from scratch using overlapping sequencing reads without a reference, which is crucial for studying novel genomes, uncovering structural variations, and exploring genetic diversity [1]. The advancement of long-read sequencing technologies, such as PacBio, Oxford Nanopore Technologies and BGI CycloneSEQ [2] platforms, has significantly improved *de novo* assembly by generating reads spanning repetitive regions and complex structural variants [3, 4]. Most long-read *de novo* assembly methods rely on the Overlap-Layout-Consensus (OLC) approach, which aligns reads based on overlaps and iteratively refines contigs to reconstruct high-quality genomes [5, 6, 7]. A key step in the OLC approach is overlap detection, which involves finding overlaps from a large collection of sequencing reads, where the existence of an overlap between a pair of reads is typically determined by their sequence similarity. Overlap detection faces two primary challenges: the computational burden of processing large amount of sequences (millions of reads for human-sized genomes), which demands efficient algorithms and substantial memory resources, and the inherent complexity caused by repetitive genomic sequences that create ambiguous overlaps [8]. These repeats often lead to fragmented or erroneous connections in the overlap graph, significantly complicating the assembly process and potentially introducing misassemblies. This issue is further exacerbated by sequencing errors, which introduce spurious overlaps and masking true overlaps. Overcoming these obstacles is crucial for accurate genome reconstruction.

Existing methods for overlap detection can be roughly categorized into two classes: seed-and-extension and locality-sensitive hashing (LSH) methods. The seed-and-extension strategy first identifies small exact or near-exact matches (seeds) between the query sequence and the target sequences. Seed matches are then filtered based on local density or other criteria. For each retained seed, the algorithm attempts to extend the alignment in both directions using dynamic programming (DP) to build a more comprehensive alignment. Finally, target sequences with the best alignments (according to certain heuristic criteria) are identified as overlapping sequences. Overlap detection based on seed-and-extension is widely adopted by both general-purpose sequence alignment tools such as BLAST [9], DALIGN [10], and minimap2 [11], as well as *de novo* assembly-oriented tools such as MECAT [12], wtdbg2 [13], PECAT [7], and xRead [14].

As seed-and-extension methods only perform time-consuming DP-based alignment on high-potential regions that contain many seed matches, they scale well to large databases and long query sequences. However, the seed matching and filtering steps are heuristic in their nature, and are prone to returning target sequences that are locally but not globally similar to the query sequence (false positives) [15]. In addition, these methods often depend on many parameters such as minimum seed density, which require careful tuning to achieve a good trade-off between accuracy and computational efficiency.

LSH-based methods, in particular MinHash, have also been widely used in overlap detection. MinHash was originally developed for text mining tasks, e.g. finding similar news articles [16] In MinHash, each sequence is first encoded as an unordered set of unique tokens, which are typically k-mers (subsequences of fixed-length *k*). The MinHash algorithm then applies multiple independent hash functions and records the minimum hash value of each set under each hash function. The resulting “sketch” serves as a compact approximation of the full set. Overlapping sequences will have more similar sketches compared to unrelated sequences. Popular *denovoassembly* tools that implement variants of MinHash include MHAP [17], Canu [5], and Shasta [6]. In addition, the overlap detection tool BLEND [18] uses another LSH-based method named SimHash to generate sketches for input sequences.

The LSH strategy replaces the time-consuming pairwise alignment step with rapid sketch comparison and therefore can potentially scale to large genomes while maintaining computational efficiency. For example, Shasta [6] is able to perform *denovo* assembly of the human genome within 6 hours, of which only 3% is spent on the MinHash step. However, a key limitation of this strategy is that highly repetitive tokens, such as k-mers from low-complexity genomic regions, can dominate the sketch, overshadowing more informative, unique tokens, affecting assembly quality in these regions [5]. Another challenge of this strategy is the difficulty in identifying overlaps between sequences that vary greatly in their lengths using their respective sketches [19, 6, 20].

To address the limitations of existing approaches for overlap detection, we developed a new strategy inspired by common single-cell sequencing workflows. Our approach, hereby referred to as FEDRANN, comprises three main steps: feature extraction (FE), dimensionality reduction (DR), and approximate nearest neighbor (ANN) search. We first designed a benchmarking pipeline to evaluate overlap graph quality using both simulated and real long-read sequencing data. Using this pipeline, we systematically compared various combinations of feature extraction, DR, and ANN methods across datasets of differing sizes and complexities. We then implemented the best-performing method, which involved inverse document frequency (IDF) transformation, sparse random projection (SRP) and ANN searching using NNdescent, as a integrated overlap detection tool named Fedrann. We benchmarked Fedrann against established tools including minimap2, MHAP, MECAT2, wtdbg2, xRead and BLEND. Our results show that Fedrann was able to construct high-quality overlap graphs for genomes of varying size and complexity without consuming excessive amount of time, highlighting its potential utility in *de novo* genome assembly.

## Results

### Overview of the FEDRANN workflow

Many text-mining algorithms, such as MinHash and SimHash [21], employ the Bag-of-Words (BoW) model, which discards the ordering of tokens within a sequence and focuses solely on the presence or absence of each token, effectively encoding the sequence database as a high-dimensional sparse matrix [22]. We observed that, once sequences are encoded in this manner, overlap detection becomes analogous to performing a nearest-neighbor search in single-cell sequencing data, where both tasks rely on identifying similar items based on some similarity or distance metric in a high-dimensional sparse space (i.e., the sequence *×* token matrix or the cell *×* gene matrix). Based on this analogy, we hypothesized that dimensionality reduction and approximate nearest-neighbor (ANN) methods, which are widely used in single-cell data analysis, could also be applied to overlap detection with appropriate adaptations. Accordingly, we developed a new approach for overlap detection (**Figure 1**) consisting of three main steps: feature extraction, dimensionality reduction (DR), and ANN search, as described below.

**Figure 1.**
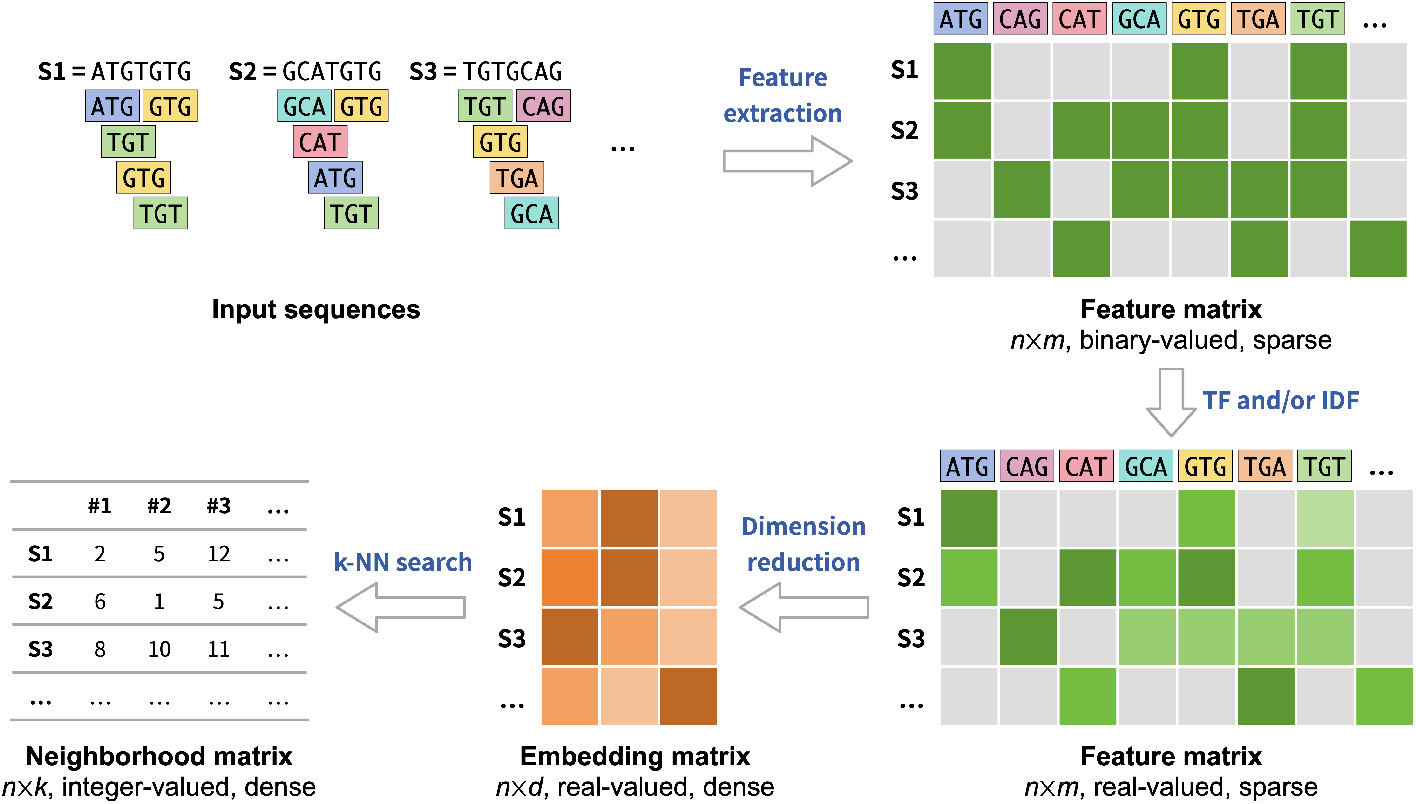
Diagrammatic representation of the proposed algorithm for sequence similarity search. The algorithm is comprised of three key steps: (1) encoding the input sequences into a high-dimensional sparse feature matrix, with optional TF and/or IDF weighting; (3) applying a dimensionality reduction technique to embed the sequences into a space of lower dimensionality; (4) identifying sequence similarity through the use of a k-nearest neighbor search algorithm.

#### Feature extraction

Given a dataset of *n* sequences, we uniformly sampled an alphabet of *m* k-mers and encoded the dataset as a high-dimensional matrix *X* with *n* rows and *m* columns, where each entry *X*[*i, j*] represents the weight of *k*-mer *j* in sequence *i*. By default, the weights were binary, indicating the presence or absence of a *k*-mer. Optionally, term frequency (TF) and/or inverse document frequency (IDF) transformations could be applied to adjust the weight of each *k*-mer based on its frequency across the dataset. Since each sequence typically contains only a small subset of all possible *k*-mers, the resulting matrix is sparse.

#### Dimensionality reduction

Although it is possible to directly apply nearest-neighbor algorithms to the high-dimensional feature matrix, doing so is often computationally expensive due to the large number of dimensions. To mitigate this, we applied dimensionality reduction techniques to project each high-dimensional feature vector into a lower-dimensional space, while aiming to preserve the relative distances between sequences.

#### ANN search

Since exact *k*-nearest neighbor (k-NN) search has quadratic time complexity, it becomes infeasible for large genomic sequencing datasets with millions of reads. To address this, we employed approximate k-NN algorithms to identify the most similar sequences for each query sequence in the dataset. These candidate overlaps were then used to define edges in the resulting overlap graph.

Many overlap detection tools, such as minimap2 and MHAP, aim to identify all pairs of overlapping reads in a dataset. Instead, FEDRANN employs a *k*-nearest neighbors (k-NN) search to retrieve only the top *k* most similar reads for each query read. These nearest neighbors are expected to exhibit the longest overlaps with the query. This approach aligns with the concept of the *best overlap graph* (BOG) [23, 5], which is grounded in the observation that full enumeration of overlapping read pairs is unnecessary, as shorter overlaps are typically redundant and pruned during graph refinement. For instance, the Shasta assembler retains by default only the top six edges per vertex in its overlap graph [6]. Restricting the number of edges in the overlap graph is advantageous, as it significantly reduces the complexity of subsequent graph layout computations.

To evaluate the performance of specific combinations of feature extraction, dimensionality reduction, and *k*-nearest neighbors (k-NN) search methods, we constructed overlap graphs for multiple long-read sequencing datasets and compared them against reference graphs derived from corresponding reference genomes (**Figure 2**). The datasets included sequencing reads from Oxford Nanopore Technologies (ONT), PacBio HiFi, and CycloneSEQ, all sampled from three repeat-rich regions of the human genome: the *HLA* immunogene cluster, the *IGK* immunoglobulin κ-light chain locus, and chromosome 22 (**Table S1**). The HLA and IGK regions each contain over 40% repetitive elements and are of notable medical relevance [24], while chromosome 22 encompasses large heterochromatic segments. For each method combination, performance was quantified by the error rate of the resulting overlap graph, defined as the proportion of incorrect edges among all edges.

**Figure 2.**
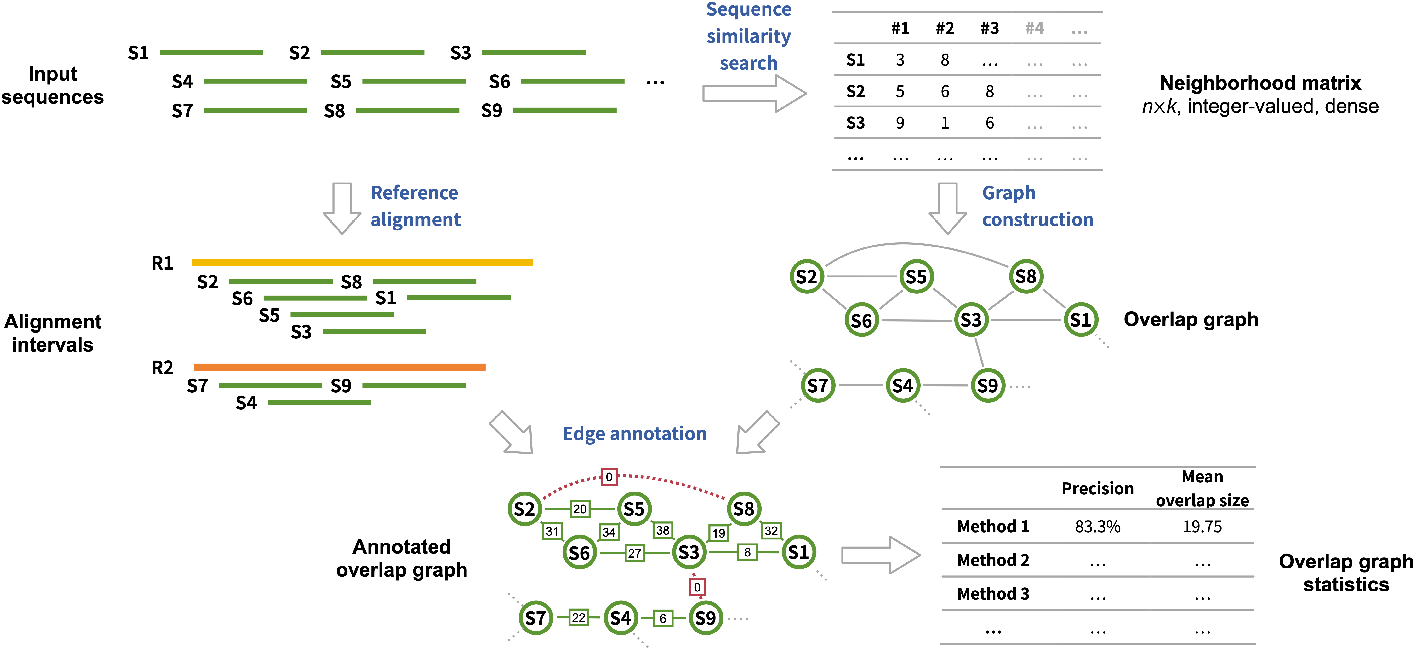
Assessment of sequence similarity search algorithms. For the analysis of each algorithm, an overlap graph is constructed by linking each query sequence to the top *k* sequences identified as most similar by the algorithm. This overlap graph is subsequently assessed against a reference graph, which is obtained by aligning the input sequences to a reference genome to ascertain the accuracy and overlap size of each edge. The algorithm’s performance is evaluated based on precision and mean overlap size.

### Inverse document frequency (IDF) transformation improves overlap detection accuracy

We first evaluated the impact of different weighting schemes (TF-IDF, TF, IDF and raw TF) during feature extraction, and compared similarity metrics (Euclidean distance and cosine distance) used in k-NN search. As the sizes of the three benchmarking regions were relatively small (3.92–5.75 Mb), we were able to use k-d tree based exact k-NN search without dimensionality reduction to inspect the upper limit of the FEDRANN strategy.

Our results showed that using cosine distance for ENN search resulted in significantly higher accuracy compared to Euclidean distance (**Figures 3, S1**, and **S2**). This is likely because Euclidean distance is sensitive to vector magnitude, while cosine distance captures only the angular difference between vectors, making it more robust to variations in sequence length. As a result, overlapping sequences of different lengths may still exhibit low cosine distances but high Euclidean distances, leading to better overlap detection when cosine similarity is used.

**Figure 3.**
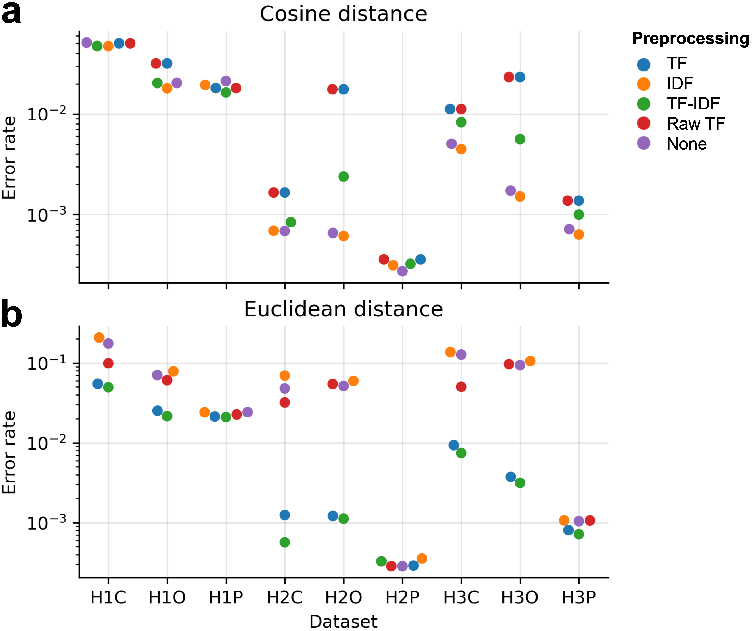
Assessment of preprocessing methods and similarity metric for overlap detection. **(a-b)** Overlap detection error rate of pipelines combining various preprocessing methods for feature extraction (TF, IDF, TF-IDF, Raw TF, and no preprocessing) and exact k-NN search based on cosine **(a)** or Euclidean **(b)** distance. Top six neighbors according to each distance metric were used to construct overlap graphs for evaluation. Abbreviations: TF, term frequency; IDF, inverse document frequency.

We evaluated the impact of different weighting schemes (TF, IDF, IDF and raw TF) on overlap graph quality. For baseline comparison, we also constructed a binary feature matrix that only records the presence (1) or absence (0) of k-mers in each read, discarding all frequency and weighting information. While TF-IDF has been used in assemblers like Canu [5] to help resolve repetitive sequences, our experiments indicated that IDF alone, rather than TF or TF-IDF, was more effective in improving graph quality. When cosine distance was used, applying IDF weighting led to better performance than both TF-IDF and unweighted representations, whereas TF alone and raw TF degraded performance relative to the unweighted baseline (**Figures 3, S1**, and **S2**). These findings suggest that high-frequency *k*-mers—such as those originating from transposable elements, short tandem repeats, or other repetitive regions—are a major source of error in overlap graph construction. By down-weighting these features, IDF enhances the ability to distinguish between true overlaps. In contrast, TF and raw TF emphasizes frequent features and may counteract the benefits of IDF, thereby reducing accuracy.

We plotted the overlap graph error rate as a function of the number of overlap candidates (*k*). As expected, the error rate increased with larger values of *k*, reflecting the greater difficulty in identifying larger numbers of correct overlaps. Notably, the combination of IDF transformation and cosine distance consistently outperformed other method combinations in terms of accuracy. Based on this observation, we selected the IDF–cosine distance combination for subsequent analyses, including evaluating the effects of read length, coverage depth, and sequencing accuracy, as well as for selecting dimensionality reduction and ANN search methods, as described below.

### Longer reads, deeper coverage, and higher accuracy improve overlap detection

We systematically evaluated the impact of read length, coverage depth, and sequencing accuracy to overlap detection by generating simulated datasets with PBSIM3 for human chromosome 22, covering read lengths of 10–30 kb, accuracy of 91%–99%, and depths of 10×–50×. We found that longer reads and deeper coverage had a large positive effect on overlap graph accuracy (**Figure S3**). We hypothesize that this was because longer reads and deeper coverage both lead to longer overlaps between adjacent reads, facilitating overlap detection. Meanwhile, improved sequencing accuracy only had a moderate positive effect on overlap graph accuracy. For example, at 30× coverage and mean read length 20 kb, increasing sequencing accuracy from 91% to 99% only reduced error rate from 8.34% to 5.55% for human chromosome 22. This observation suggested that our FEDRANN strategy is relatively robust to sequencing errors.

### Sparse random projection enables scalable and accurate dimensionality reduction

Dimensionality reduction (DR) techniques are widely used in single-cell sequencing analyses to reduce noise, accelerate computation, and facilitate data visualization. To evaluate their utility in overlap graph construction, we tested a range of DR methods, including linear approaches such as principal component analysis (PCA) [25] and sparse random projection (SRP), as well as non-linear methods including Uniform Manifold Approximation and Projection (UMAP) [26], Spectral Embedding, and scBimapping. We also evaluated SimHash [27], a locality-sensitive hashing (LSH)–based method related to random projection, which has been widely used in text mining and recently applied to biological sequence analysis [18, 28].

For each method, we first constructed the input feature matrix using IDF weighting, then applied the DR method, and finally computed exact *k*-nearest neighbors (ENN) from the resulting low-dimensional embeddings (**Figure 1**). We found that all tested DR methods except UMAP were able to generate accurate overlap graphs for smaller datasets (**Figure 4a**), suggesting that they preserved the relative pairwise distances between sequences sufficiently well. However, when applied to larger datasets, most methods failed to complete within the predefined resource constraints (wall clock time *≤* 6 hours; peak memory *≤* 1 TB) (UMAP), or the matrix size surpassed the maximum constraints of the methods (PCA, Spectral and scBiMapping), resulting in dimensionality reduction failure. The only methods that successfully scaled to the largest datasets, H3C, H3O and H3P, was SRP, GRP and SimHash. Time and memory profiling on dataset H3P further confirmed that SRP was the fastest method and had the lowest memory usage among those tested (**Figure 4b**).

**Figure 4.**
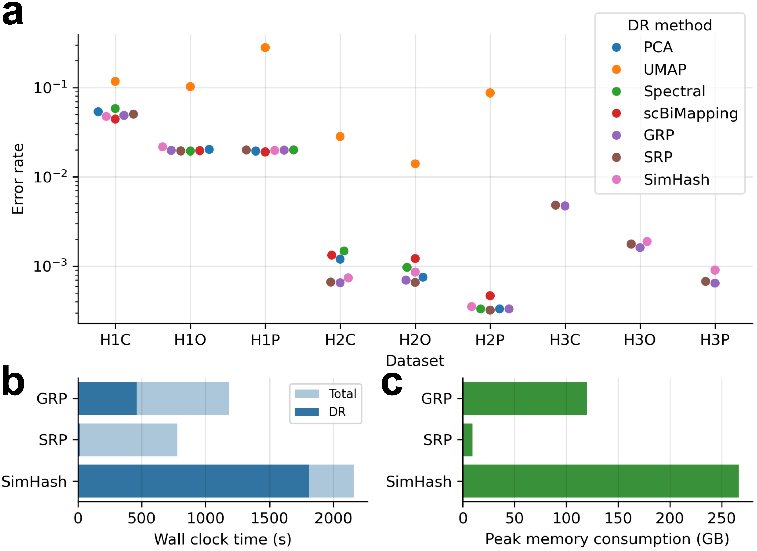
Assessment of dimensionality reduction methods. (a) Overlap detection error rate of various dimensionality reduction methods. IDF preprocessing was used in feature extraction. Exact Hamming distance (for SimHash) or cosine distance (for other methods) was used in k-NN search. Top six neighbors according to each distance metric were used to construct overlap graphs for evaluation. Missing dots indicate that the corresponding methods failed to generate results under predefined computational constraints (wall clock time *≤* 6 h and peak memory *≤* 900 GB). **(b-c)** Wall clock time **(b)** and peak memory consumption **(c)** of various dimensionality reduction methods in dataset H3P. Methods that failed to generate results are not shown. Abbreviations: PCA, principal component analyses; UMAP, Uniform Manifold Approximation and Projection; Spectral, spectral embedding; GRP, Gaussian random projection, SRP: sparse random projection; DR, dimensionality reduction.

The accuracy of *k*-NN search after DR typically depends on the embedding dimension, as higher dimensions allow more information from the original feature matrix to be retained. To characterize this relationship, we varied the embedding dimension for each dataset using SRP. The results showed that increasing the embedding dimension improved accuracy, but with diminishing returns (**Figures S4** and **S5**). Most datasets achieve the highest precision with 1,000 dimensions, while some smaller regions require fewer dimensions to reach peak precision.

To better understand this trend, we examined the correlation between pairwise cosine distances before and after SRP in dataset H3C. The results showed a strong positive correlation that increased with embedding dimension, with the Pearson *R*^2^ reaching 0.99 at 3000 dimensions (**Figure S6**). Based on these findings, we selected SRP as the DR method for further benchmarking due to its scalability, efficiency, and strong distance-preserving properties.

### NNDescent offers the best trade-off between speed and accuracy among ANN methods

In addition to dimensionality reduction, another strategy to accelerate k-NN search is to use approximate k-NN (ANN) algorithms, which leverage indexing structures or heuristics to reduce the computational cost of neighbor retrieval. We evaluated five ANN approaches that are widely used in single-cell sequencing and information retrieval applications: NNDescent [27], Hierarchical Navigable Small World (HNSW) [29], product quantization (PQ) [30], inverted file index with product quantization (IVF-PQ) [30], and random projection forest (RPF) [31]. All ANN methods were applied to low-dimensional embeddings generated by sparse random projection (SRP).

Among these methods, NNDescent and HNSW achieved high precision when used to construct overlap graphs, whereas PQ, IVF-PQ and RPF performed less favorably in terms of accuracy (**Figure 5a**). In terms of computational efficiency, NNDescent and IVF-PQ were the fastest, while RPF consumed the least memory (**Figure 5b**). Across all configurations, the ANN step emerged as the computational bottleneck in the FEDRANN pipeline, often requiring significantly more time than the SRP step (**Figure 5b**), although both steps had comparable peak memory usage (**Figures ??b** and **5b**). Therefore, we found NNDescent to be the most useful ANN method for our workflow. It offered the best trade-off between speed and accuracy, making it well-suited for scalable overlap graph construction when combined with SRP-based dimensionality reduction.

**Figure 5.**
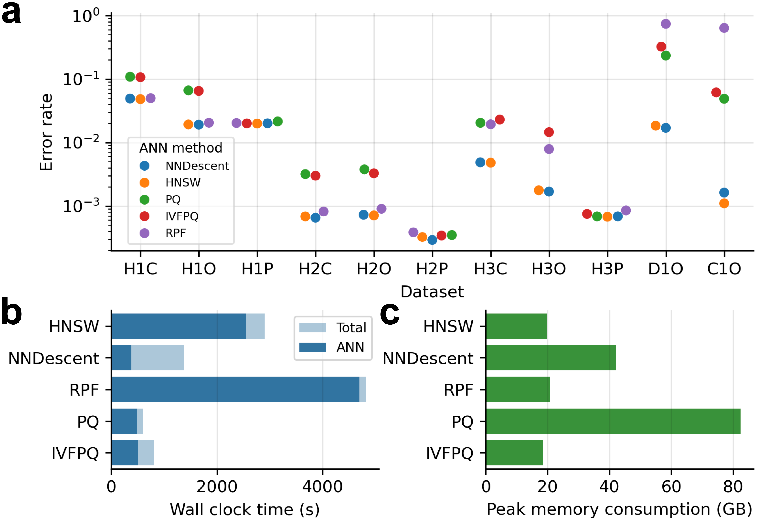
Assessment of ANN methods. **(a)** Overlap detection error rate of various ANN methods. IDF preprocessing was used in feature extraction. Feature matrices were reduced to 1,000 dimensions using Sparse Random Projection prior to ANN search. Cosine distance was used in ANN search. Top six neighbors were used to construct overlap graphs for evaluation. Missing dots indicate that the corresponding methods failed to generate results under predefined computational constraints (wall clock time *≤* 6 h and peak memory *≤* 900 GB). **(b-c)** Wall clock time **(b)** and peak memory consumption **(c)** of selected ANN methods in dataset D1O. Abbreviations: ANN, approximate *k*-nearest neighbors; HNSW, Hierarchical Navigable Small World; PQ, Product Quantization; IVFPQ, Inverted File with Product Quantization; RPF, Random Projection Forest.

### Fedrann enables high-precision, time-efficient overlap detection on large genomes

We implemented the IDF-SRP-NNDescent pipeline as a standalone tool for overlap detection, named *Fedrann. Fedrann* accepts FASTQ or FASTA files as input and identifies *k* (default 20) overlap candidates for each sequence. It outputs two plain-text files: one containing overlap information (including sequence indices, distance metrics, and neighbor ranks), and another providing associated metadata (sequence names and orientations). The tool is primarily written in Python and leverages efficient sparse and dense matrix structures via `NumPy` [32] and `SciPy` [33]. A range of optimizations were applied to improve computational performance (see Methods for details). The resulting implementation supports parallel execution and takes advantage of highly optimized matrix operations, enabling application to whole-genome long-read datasets.

We benchmarked *Fedrann* against a set of established overlap detection tools: minimap2 (in all-vs-all mode), MECAT2, MHAP, BLEND, wtdbg2, and xRead. Benchmarking was performed on real whole-genome sequencing datasets from three species with differing genome sizes and complexities: *Caenorhabditis elegans, Drosophila melanogaster*, and *Homo sapiens* (**Table S1**).

A key challenge in benchmarking arose from differences in tool design. Some tools (minimap2, MECAT2, MHAP, BLEND, and wtdbg2) aim to find all overlapping pairs, typically achieving high recall. Others (xRead and *Fedrann*) return only a subset of the top overlap candidates per read, trading recall for higher precision [14]. To enable fair comparisons, we post-processed the output of all tools to retain only the top *k* candidates (with *k* = 6, 12, or 18) ranked by overlap size for each read. Three additional metrics were used to evaluate overlap graph quality: mean overlap size, the number of correct overlap candidates per read (#COC), and the number of connected components in the graph (#CC). Mean overlap size measures whether the ability to identify the longest overlaps for each read, as longer overlaps are typically more informative and can be used to infer shorter ones. Higher #COC values indicate better per-read connectivity, while lower #CC values suggest less fragmentation and higher contiguity.

A memory limit of 950 GB was enforced during benchmarking. Under this constraint, MHAP failed to complete on all three human datasets (H4C, H4O, and H4P), and xRead failed on H4C. All other tools completed successfully (**Figure 6**). *Fedrann* achieved the highest accuracy on four out of five datasets (**Figure 6a**); the exception was H4C, where MECAT2 performed slightly better. wtdbg2 also showed strong accuracy, consistently ranking second or third across datasets. In terms of mean overlap size, MECAT2 consistently ranked lowest, while other tools performed comparably (**Figure 6b**). For graph contiguity, *Fedrann*, wtdbg2, MHAP, and MECAT2 generally outperformed the other methods, with no clear single leader (**Figure 6c**). Analysis of low-connectivity reads under varying #COC thresholds showed that xRead produced a large number of poorly connected reads—for instance, in dataset H4P, over 50% of reads had fewer than five correct edges. *Fedrann* ranked first or second across all datasets.

**Figure 6.**
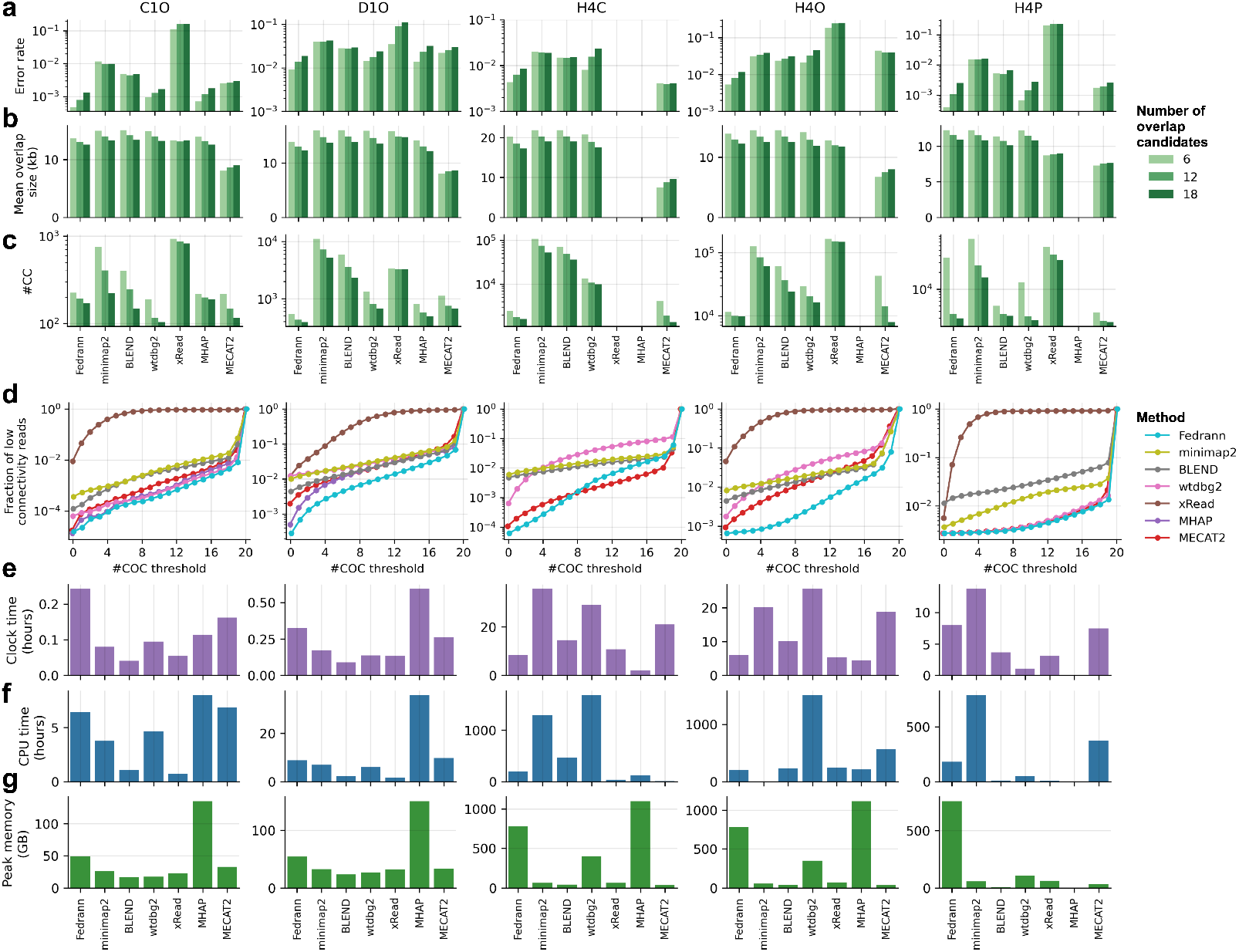
Benchmarking of overlap detection tools. **(a-c)** Overlap detection error rate **(a)**, mean overlap size **(b)**, and #CC **(c)** of various overlap detection tools in five whole-genome datasets (columns). Top 6, 12, or 18 overlap candidates (colors) according to cosine distance (for Fedrann) or overlap size (for other methods) were used to construct overlap graphs for evaluation. **(d)** Fraction of low-connectivity reads plotted against #COC threshold for various overlap detection tools. **(e-g)** Wall-clock time **(e)**, CPU time **(f)**, and peak memory **(g)** usage of various overlap detection tools. In **(a-c)** and **(e-g)**, black crosses indicate that the corresponding methods failed to generate results under predefined computational constraints (wall clock time *≤* 72 h and peak memory *≤* 900 GB). These methods were not included in **(d)**. Abbreviations: #CC, number of connected components; #COC, number of correct overlap candidates.

In terms of computational cost, all tools completed the smaller datasets (C1O and D1O) within one hour and with moderate memory usage (<60 GB), except MHAP, which consumed up to 150 GB (**Figure 6e–g**). For the larger human datasets (H4C, H4O, H4P), where computational performance is especially critical, *Fedrann*, BLEND, and xRead were among the fastest in terms of both wallclock and CPU time. Interestingly, wtdbg2 took over 20 hours for ONT and CycloneSEQ datasets (H4C and H4O), but only 1.1 hours for the HiFi dataset (H4P). minimap2, BLEND, xRead, and MECAT2 demonstrated low memory usage (<100 GB), whereas *Fedrann* required over 700 GB for the human genome datasets.

In summary, *Fedrann*, wtdbg2, and MECAT2 emerged as the best-performing tools when considering both graph quality and computational efficiency. *Fedrann* offered a balanced trade-off across accuracy, overlap size, graph contiguity, and speed, though it incurred a high memory footprint. wtdbg2 was accurate and memory-efficient but lagged in contiguity, connectivity, and speed (except for HiFi data). MECAT2 was both accurate and efficient in memory usage, but failed to capture some of the longest overlaps. Overall, our results highlight *Fedrann* as a robust and competitive new tool for long-read overlap detection.

## Discussion

We proposed a novel strategy for overlap graph construction that integrates dimensionality reduction (DR) and approximate nearest neighbor (ANN) techniques. We conducted comprehensive evaluations on both simulated and real long-read sequencing datasets. Among the tested combinations, the pipeline consisting of inverse document frequency (IDF) transformation, sparse random projection (SRP), and NNDescent achieved the best trade-off between accuracy and computational efficiency. Based on these findings, we developed an efficient, open-source, and userfriendly implementation named Fedrann. Benchmarking results showed that Fedrann generated overlap graphs that were comparable to or better than those produced by state-of-the-art tools such as MECAT2 and wtdbg2, in terms of both overlap accuracy and contiguity for large genomes, without sacrificing computational speed.

Although the concept of framing overlap graph construction as a special case of *k*-nearest neighbor search has been explored previously [6], to our knowledge, the explicit application of DR and ANN algorithms to this task has not been reported. Our results demonstrate the potential of the Fedrann strategy as a robust and scalable alternative for overlap detection.

The current implementation of Fedrann was written primarily in Python and relies on general-purpose numerical libraries that are not specifically optimized for nucleotide sequence processing. Despite this, Fedrann was able to construct an overlap graph for the human genome within 5–8.5 hours, comparable to other overlap detection tools implemented in compiled languages such as C and C++. This computational efficiency is likely attributable to the use of dense and sparse matrix representations for input sequence databases, enabling highly optimized operations via Python libraries with C extensions such as `NumPy`, `SciPy`, and `pyNNDescent`.

However, memory consumption remains a key limitation, particularly during the ANN search step using NNDescent, which involves maintaining multiple data copies in memory. Future work may focus on reducing the memory footprint through the use of shared memory, memory-mapped files, or other memory-efficient data structures.

Beyond computational performance, another strength of using matrix-based representations is the clarity and modularity of the pipeline. Most parameters in Fedrann have well-defined meanings and predictable effects on the output, typically reflecting trade-offs between overlap detection accuracy and resource usage, without being tightly coupled to a specific sequencing platform. For instance, increasing the embedding dimension in SRP generally improves accuracy at the cost of additional computation. This platform-agnostic design allows Fedrann to be easily adapted to emerging sequencing technologies with minimal parameter tuning.

In this study, we limited our evaluation to core overlap detection metrics such as precision, overlap size, and runtime. However, the quality of the final *de novo* assembly often depends on additional properties of the overlap graph that are not captured by these metrics. For example, consistent connectivity across genomic regions can significantly influence the contiguity and completeness of the assembled genome, though such properties are challenging to quantify. An important direction for future research is to incorporate the Fedrann strategy into a complete OLC-based *de novo* assembly pipeline, and to evaluate the assembled genomes in terms of standard metrics such as contiguity (e.g., NGA50), completeness (e.g., BUSCO scores), and base-level accuracy.

Finally, we note that overlap graph construction can be viewed as a special case of the broader sequence similarity search (SSS) problem, where the goal is to retrieve the most similar sequences for each query from a large target database. Our methodology could be extended to other SSS tasks with appropriate modifications. One promising application is taxonomic classification in metagenomics, where the query sequences are reads or contigs, and the target database consists of pre-indexed microbial reference genomes. Unlike overlap detection, which operates on all sequences simultaneously, classification tasks often require fast querying against static indexes. The IDF-SRP-NNDescent pipeline is naturally suited for such use cases: both IDF and SRP are linear and independent transformations, and NNDescent supports incremental querying. Nevertheless, further empirical validation is necessary to assess the performance of DR and ANN techniques in metagenomic classification and related sequence analysis problems.

## Methods

### Fedrann implementation

Fedrann was developed as a command-line tool designed to identify candidate overlapping sequence pairs from an input FASTA or FASTQ file containing multiple sequences based on the FEDRANN workflow. The tool is primarily implemented in Python 3 and executes in three stages, as detailed below:

#### Feature extraction

We first employed the high-performance k-mer counting tool JellyFish [34] to extract k-mers with multiplicities exceeding a predefined threshold, thereby filtering out low-frequency k-mers likely arising from sequencing errors. A k-mer alphabet was constructed by uniformly sampling a predefined fraction (default: 10%) from the set of extracted k-mers. We then used a custom multi-threaded C++ program to determine the presence of each kmer in the k-mer alphabet across all input sequences (and their reverse complements). This information was used to construct a sparse feature matrix, where rows correspond to sequences and columns to k-mers, as previously described. The feature matrix was stored in memory using the Compressed Sparse Row (CSR) format. Associated metadata, including read names and orientations, were recorded and saved to disk.

An inverse document frequency (IDF) transformation was then applied to the feature matrix using efficient sparse matrix operations.

#### Dimensionality reduction

We implemented a custom, multi-process version of Sparse Random Projection (SRP) to reduce the dimensionality of the feature matrix. To minimize memory overhead and avoid redundant data copying, our implementation leverages shared memory for efficient parallelization of the matrix multiplication step. A random sparse projection matrix is generated once in the main process and shared with all child processes. The input matrix is split into chunks (each with 100,000 rows by default), and each process independently projects its assigned chunk. The resulting embeddings from all processes are written into a shared dense output array for downstream use.

##### Approximate nearest neighbor search

We used the PyNNDescent library to efficiently compute neighborhood relationships from the embedding matrix in a highly parallelized manner. The resulting neighborhood graph, along with the associated similarity scores, was then converted into a candidate overlap table, which was saved to disk as the final output.

### Sequencing datasets

To evaluate various overlap detection methods, we used genomic sequencing data from three model species: *Homo sapiens, Caenorhabditis elegans*, and *Drosophila melanogaster*. The corresponding reference genomes were obtained from the following sources: the complete human T2T-CHM13 assembly v2.0 [3], the *Caenorhabditis elegans* WBcel235 assembly (WormBase release WS285; NCBI Assembly ID: GCA_000002985.3),and the *Drosophila melanogaster* Release 6 genome assembly (NCBI Assembly ID: GCF_000001215.4).

Simulated sequencing datasets with varying read lengths, error rates, and coverage depths were generated using PBSIM3 [35], based on the `ERRHMM-ONT-HQ` model. Real sequencing datasets from Oxford Nanopore Technologies (ONT), and PacBio HiFi platforms were obtained from publicly available databases (**Table S1**). ONT R10 sequencing data for HG002 was downloaded from https://epi2me.nanoporetech.com/gm24385_2020.11/. ONT R9 sequencing data for *C. elegans* (SRR10028111) and *D. melanogaster* (SRR13070625) were retrieved from the NCBI Sequence Read Archive (SRA). PacBio HiFi sequencing data for the HG002 sample was downloaded from the Genome in a Bottle (GIAB) project [36] (https://ftptrace.ncbi.nlm.nih.gov/ReferenceSamples/giab/data_indexes/AshkenazimTrio/sequence.index.AJtrio_PacBio_CCS_15kb_10022018.HG002). CycloneSEQ G400-ER sequencing data for HG002 was generated in-house using genomic DNA extracted from the HG002 lymphoid cell line following standard library preparation and sequencing protocols.

Three genomic regions from the human genome were used for selecting appropriate methods for feature extraction, dimensionality reduction and k-NN search (**Table S1**), including the *HLA* immunogene cluster, the *IGK* immunoglobulin κ-light chain locus, and chromosome 22 (**Table S1**). For each region, real sequencing data were mapped to the corresponding reference genome using minimap2 with the following parameters: `-k 19 -w 5 -A 3 -B 2 -m 250 --secondary=no`, and reads belonging to the given region were extracted. Whole-genome sequencing data for *C. elegans* (ONT R10), *D. melanogaster* (ONT R10) and *H. sapiens* (ONT R10, PacBio HiFi, and CycloneSEQ G400-ER) were used for benchmarking overlap detection tools.

### Overlap graph construction and evaluation

For a dataset of *n* sequencing reads, an undirected overlap graph *G* = (*V, E*) was constructed from pairs of reads identified as overlapping. Each vertex in the graph represented an oriented read—either in its forward or reverse-complement orientation— resulting in a total of 2*n* vertices. An edge {*u, v*} was added to the graph if read *u* was among the *k* nearest neighbors of read *v*, or vice versa, indicating that the two reads likely originated from overlapping genomic regions. The total number of edges in the graph ranged from *kn* to 2*kn*, depending on the degree of mutuality among nearest-neighbor relationships (i.e., whether overlaps were reciprocal or one-sided).

For each benchmarking dataset, overlap graphs constructed using nearest neighbors identified by a specific method were evaluated against a reference graph *G′* = (*V, E′*) built from the same dataset. The reference graph shared the same set of vertices *V* as the overlap graphs. For real sequencing data, the reference edges *E′* were defined based on the alignment positions of reads in the reference genome. Sequencing reads were filtered based on the following criteria to remove ambiguously aligned reads: (1) read length *≥* 5 kb; (2) aligned fraction *≥* 50%; (3) mapping quality *≥* 30. For simulated data, the edges *E′* were determined according to the genomic intervals from which each read was simulated, as reported by PBSIM3.

For each edge {*u, v*} identified in overlap graph *G* = (*V, E*), if the same edge exist in the reference graph *G′*, this edge was considered correct. Otherwise, this edge was considered incorrect. We used four metrics to quantitatively evaluate the quality of an overlap graph: error rate, mean overlap size, the number of correct overlap candidates (#COC) and the number of connected components (#CC). The error rate was defined as the number of incorrect edges divided by the number of total edges. Mean overlap size was defined as the arithmetic mean of overlap size of all edges. The overlap size of incorrect edges were considered zero for this calculation. #COC was defined as the degree of each node after removing any incorrect edges. If a node does not have any correct edges, it was referred to as a singleton. #CC was defined as the number of connected components of the overlap graph after removing any incorrect edges.

### Benchmarking overlap detection tools

We benchmarked Fedrann (v0.4.27) against six state-of-the-art tools: minimap2 (v2.24), xRead (v1.0.0), BLEND (v1.0.0), MHAP (v2.1.1), MECAT2 (v20190314), and wtdbg2 (v2.5). All alignments were filtered using two criteria: (1) *≥* 100 matched bases and (2) *≥* 10% alignment identity. Notably, for minimap2 PAF files (which produced exceptionally large outputs exceeding 4TB for human whole-genome sequencing read alignments), we increased the identity threshold to 30% to reduce computational overhead. For each read, top *k* longest overlaps were selected for overlap graph construction. Refer to **Table S4** for the specific parameters used for each tool.

## Data and code availability

The CycloneSEQ G400-ER sequencing data generated in this study is available via the publicly available database CNGBdb (accession number XXXXXX). The source code for Fedrann is released under the GNU General Public License v3.0 via GitHub (https://github.com/jzhang-dev/FEDRANN).

## Ethics statement

This study was conducted in adherence to ethical standards to ensure the responsible and respectful treatment of all data and materials. Ethical approval for this study was obtained from the Institutional Review Board of BGI (FT 20060, FT 17099).

## Acknowledgments

This work was supported by National Key R&D Program of China (No.2024YFC3406300) and “Pioneer” and “Leading Goose” R&D Program of Zhejiang (No.2024C03004).

## Supplementary information

### Supplementary figures

**Figure S1.**
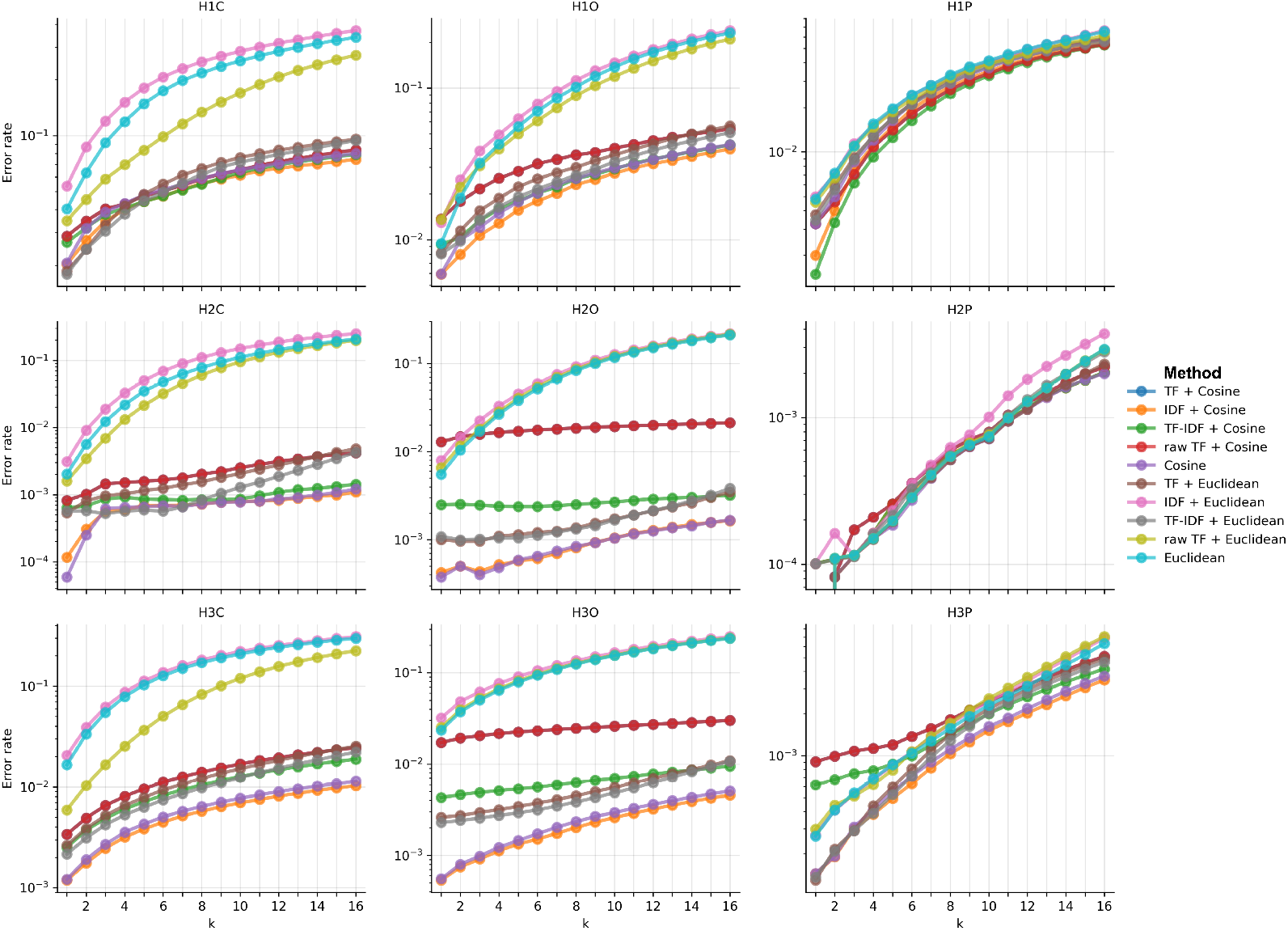
Error rate of preprocessing methods and distance metrics for overlap dectection. Overlap detection error rate of various preprocessing methods combined with Euclidean or cosine distance in various datasets. Each dot represents top-k neighbors error rate. No dimensionality reduction or ANN methods were used in k-NN search.

**Figure S2.**
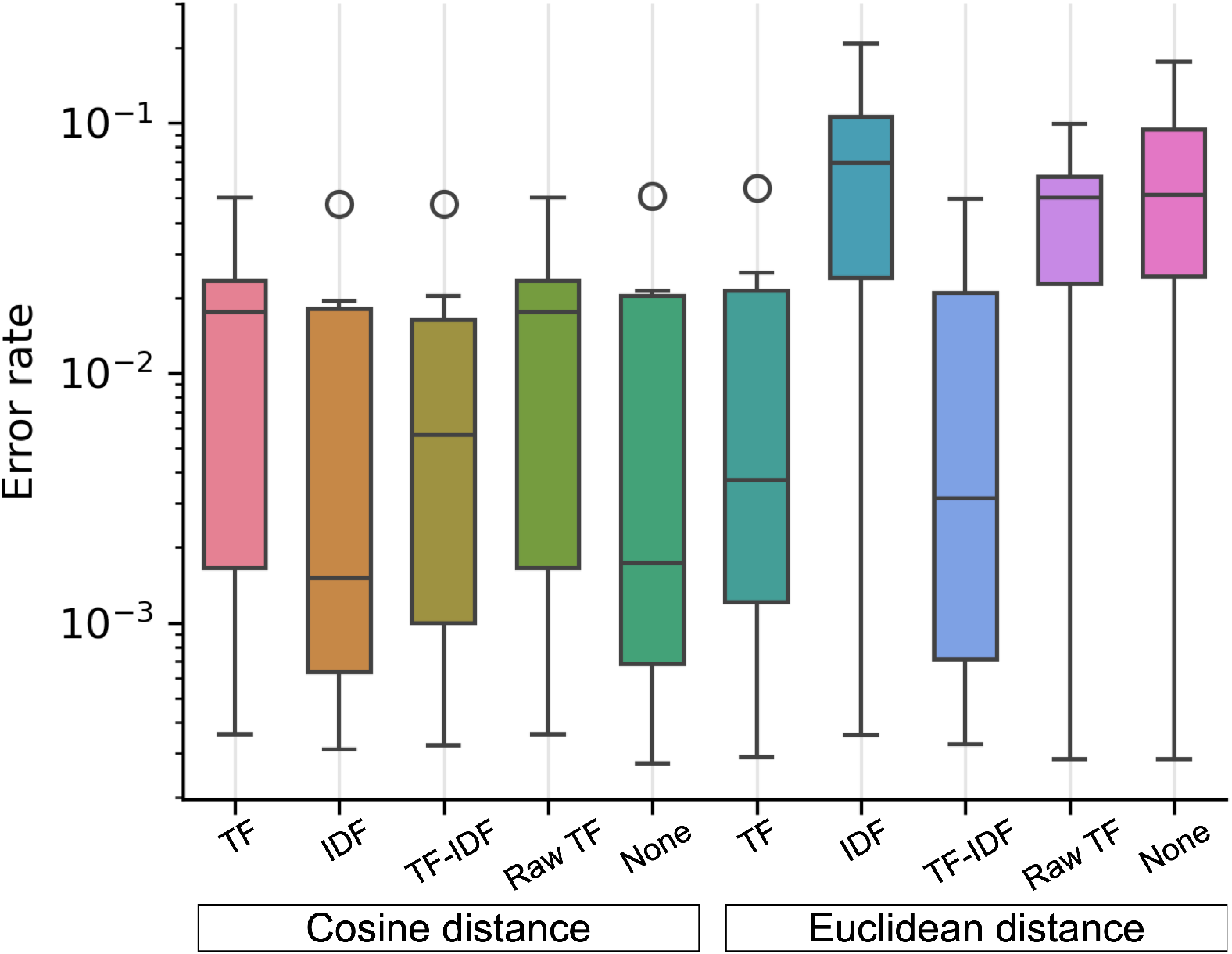
Error rate distribution of preprocessing methods and distance metrics for overlap dectection. Overlap detection error rate distribution in overlap detection across datasets (H1C-H3C, H1O-H3O, H1P-H3P) using different text preprocessing methods (TF, IDF, TF-IDF, Raw TF, None) with cosine and Euclidean distance metrics. No dimensionality reduction or ANN methods were used in k-NN search. Boxplots show quartile ranges with whiskers indicating 1.5×IQR, and circles denote outliers.

**Figure S3.**
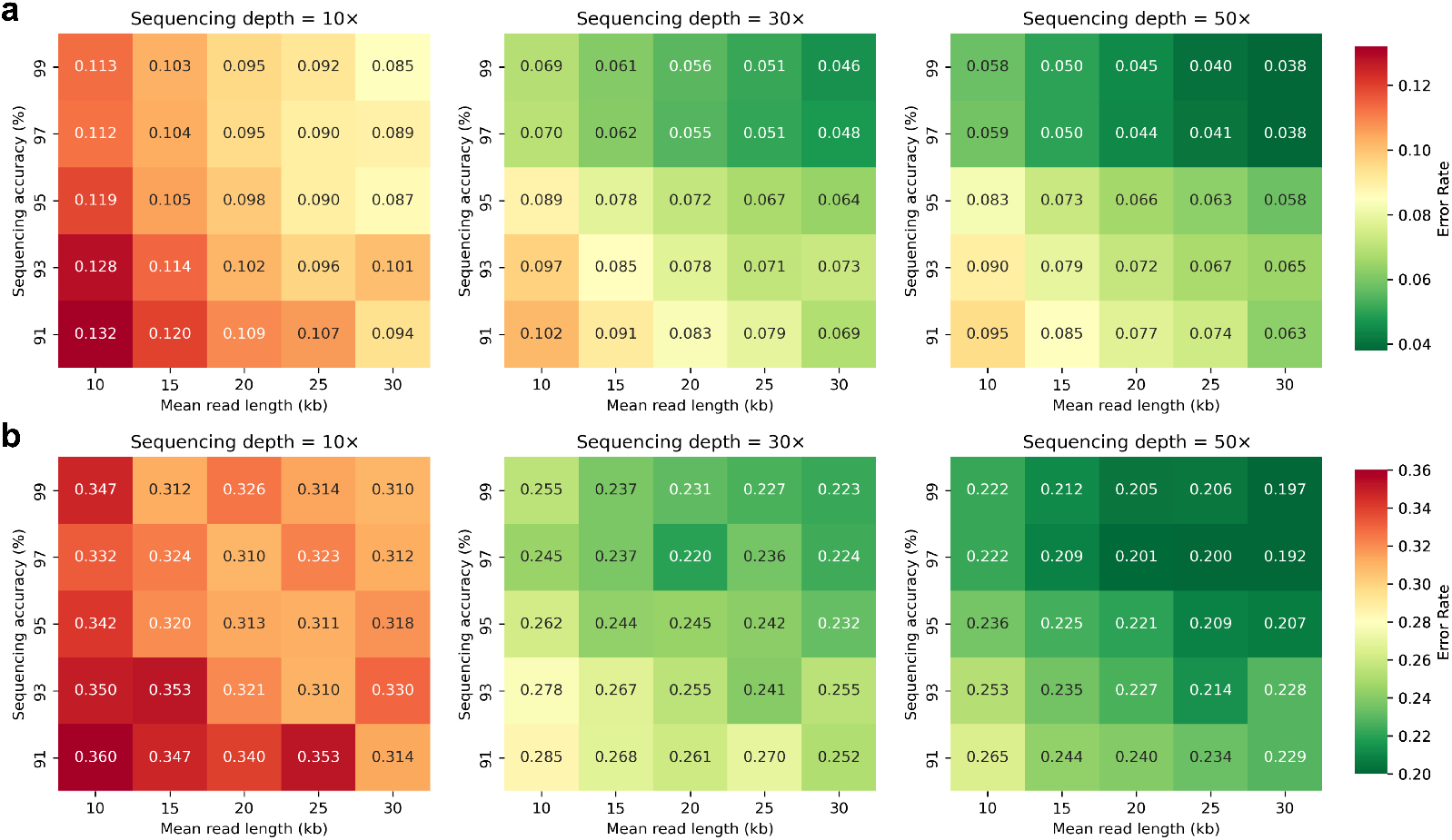
Assessment of simulated sequence reads. **(a-b)** Heatmap showing overlap detection error rates for simulated sequencing reads from the human IGK region **(a)** and chromosome 22 **(b)**. Each cell representing error rates under different combinations of sequencing depth (10×, 30×, 50×), accuracy (91%–99%), and read length (10–30 kb). IDF preprocessing was used in feature extraction. Exact cosine distance was used in k-NN search. Top six neighbors according to each distance metric were used to construct overlap graphs for evaluation. The heatmap used a red-to-green color gradient, where red indicates higher error rates and green indicates lower error rates.

**Figure S4.**
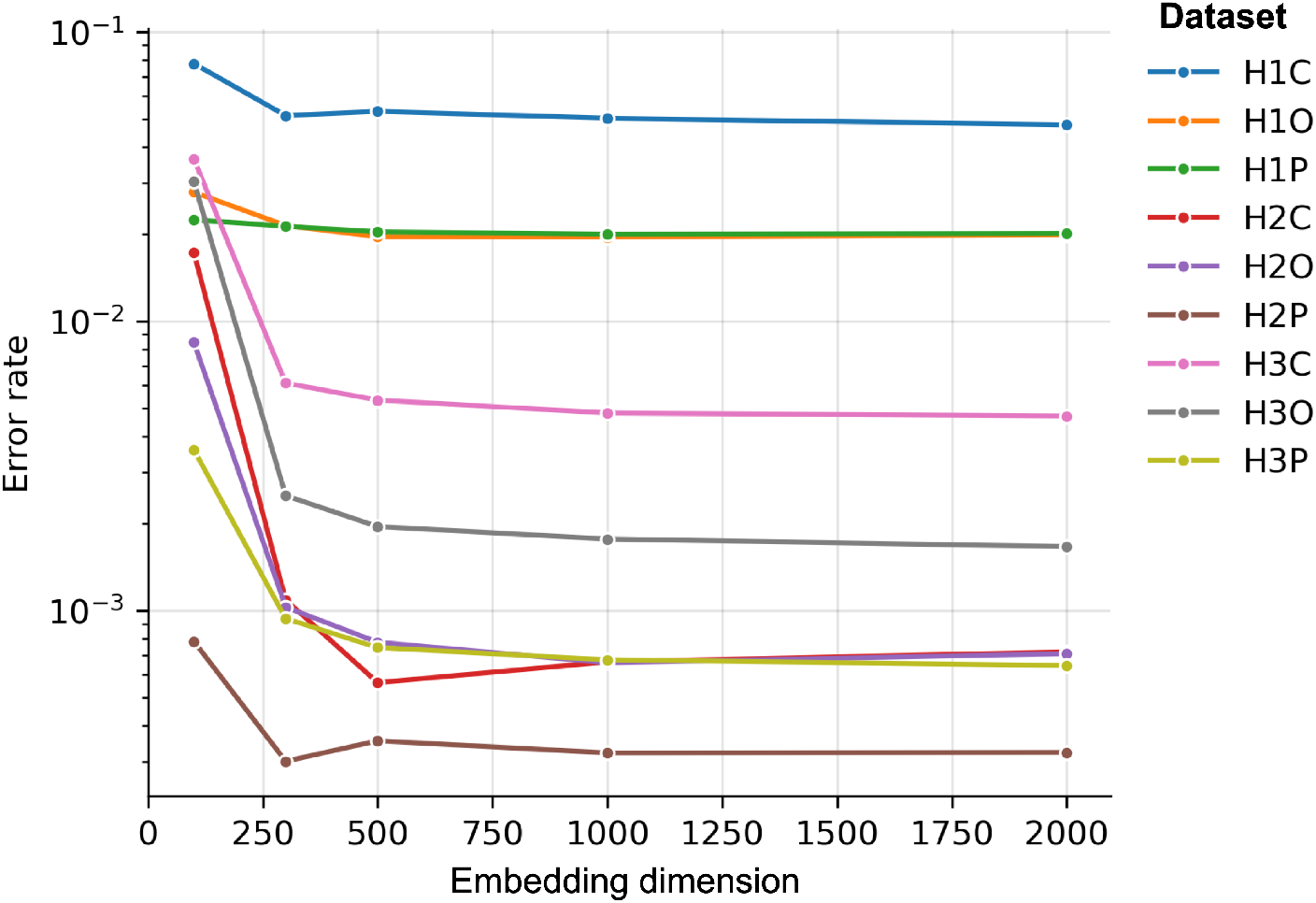
Assessment of embedding dimensions. Overlap detection error rate of various embedding dimensions. IDF preprocessing was used in feature extraction. Cosine distance was used as metric in k-NN search. Feature matrices were reduced to different dimensions using Sparse Random Projection prior to ENN search. Top six neighbors were used to construct overlap graphs for evaluation.

**Figure S5.**
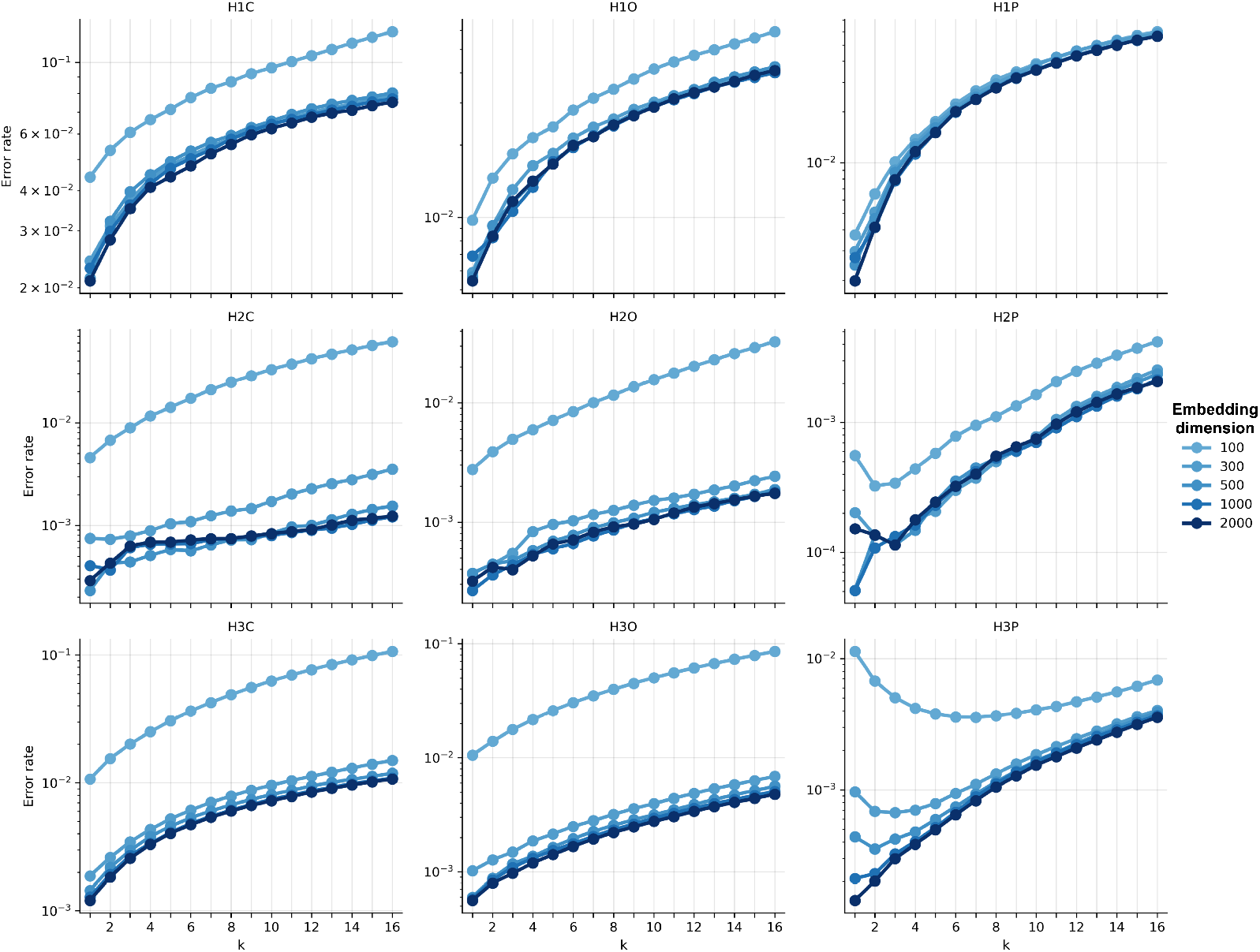
Assessment of embedding dimensions (Detail). Overlap detection error rate of various embedding dimensions. IDF preprocessing was used in feature extraction. Cosine distance was used as metric in k-NN search. Feature matrices were reduced to different dimensions using Sparse Random Projection prior to ENN search. Top six neighbors were used to construct overlap graphs for evaluation.

**Figure S6.**
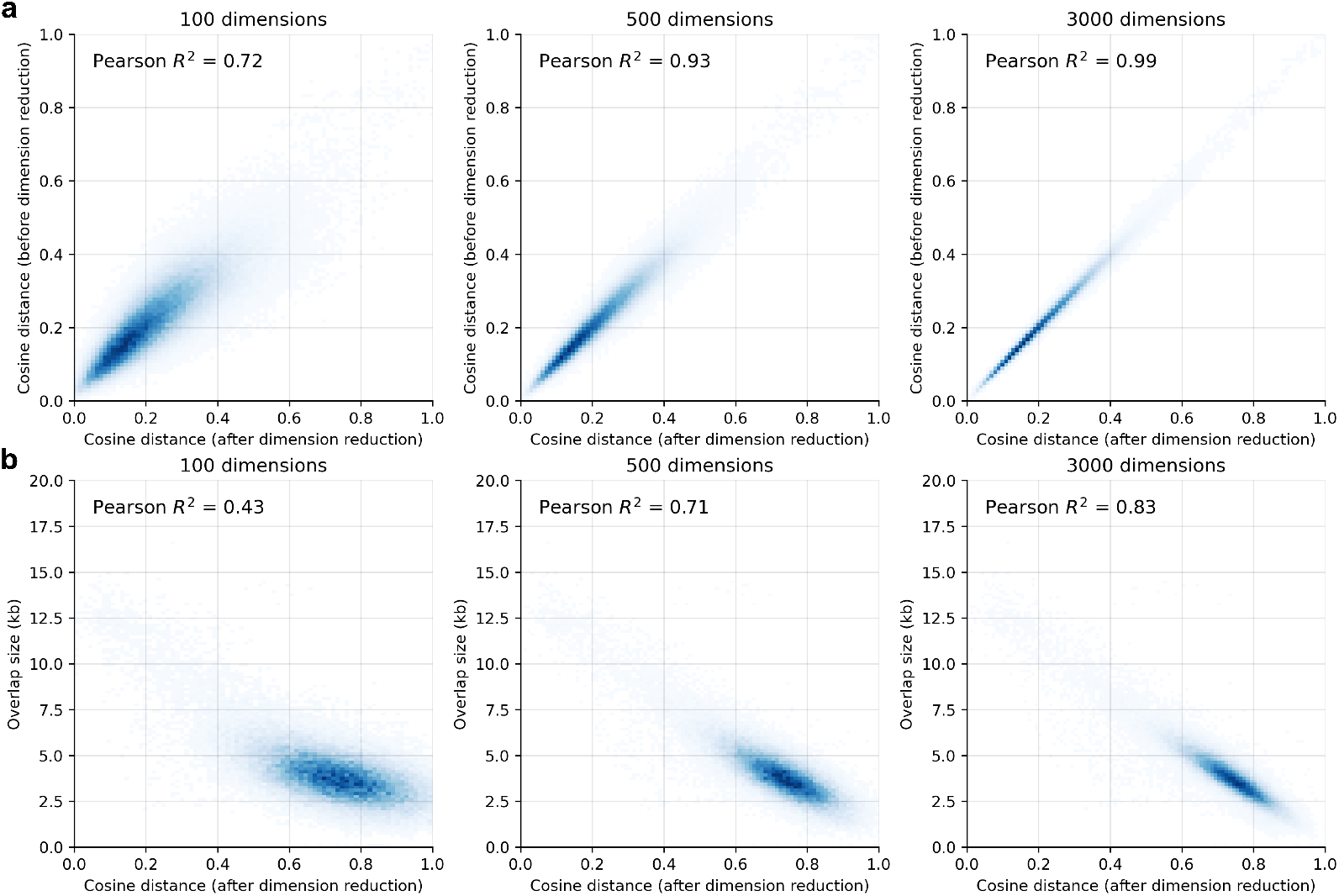
Effect of embedding dimension to distance preservation. **(a)** Correlation between distance metrics before/after dimensionality reduction at three embedding dimensions (100,500,3000) for dataset H3P. **(b)** Correlation between cosine distance (after dimensionality reduction) and overlap size at three embedding dimensions (100,500,3000) for dataset H3P.

### Supplementary tables

**Table S1.** Information of real datasets used in this study.

| ID | Sample | Region | Platform | Reference size (Mb) | Read count | N50 (kb) |
| --- | --- | --- | --- | --- | --- | --- |
| H1C | *H. sapiens* (HG002) | IGK | CycloneSEQ G400-ER | 3.92 | 2,506 | 41.91 |
| H1O | *H. sapiens* (HG002) | IGK | ONT R10 | 3.92 | 2,701 | 22.97 |
| H1P | *H. sapiens* (HG002) | IGK | PacBio HiFi | 3.92 | 2,624 | 13.48 |
| H2C | *H. sapiens* (HG002) | HLA | CycloneSEQ G400-ER | 5.75 | 11,176 | 40.67 |
| H2O | *H. sapiens* (HG002) | HLA | ONT R10 | 5.75 | 12,576 | 23.16 |
| H2P | *H. sapiens* (HG002) | HLA | PacBio HiFi | 5.75 | 12,872 | 13.42 |
| H3C | *H. sapiens* (HG002) | Chromosome 22 | CycloneSEQ G400-ER | 51.32 | 76,601 | 41.25 |
| H3O | *H. sapiens* (HG002) | Chromosome 22 | ONT R10 | 51.32 | 81,891 | 22.78 |
| H3P | *H. sapiens* (HG002) | Chromosome 22 | PacBio HiFi | 51.32 | 85,993 | 13.43 |
| H4C | *H. sapiens* (HG002) | whole genome | CycloneSEQ G400-ER | 3,117.29 | 5,127,109 | 40.67 |
| H4O | *H. sapiens* (HG002) | whole genome | ONT R10 | 3,117.29 | 6,226,490 | 23.06 |
| H4P | *H. sapiens* (HG002) | whole genome | PacBio HiFi | 3,117.29 | 6,061,187 | 13.48 |
| C1O | *C. elegans* (strain N2) | whole genome | ONT R9 | 100.29 | 221,513 | 19.02 |
| D1O | *D. melanogaster* (BDGP genome strain) | whole genome | ONT R9 | 143.73 | 281,881 | 23.92 |

**Table S2.**
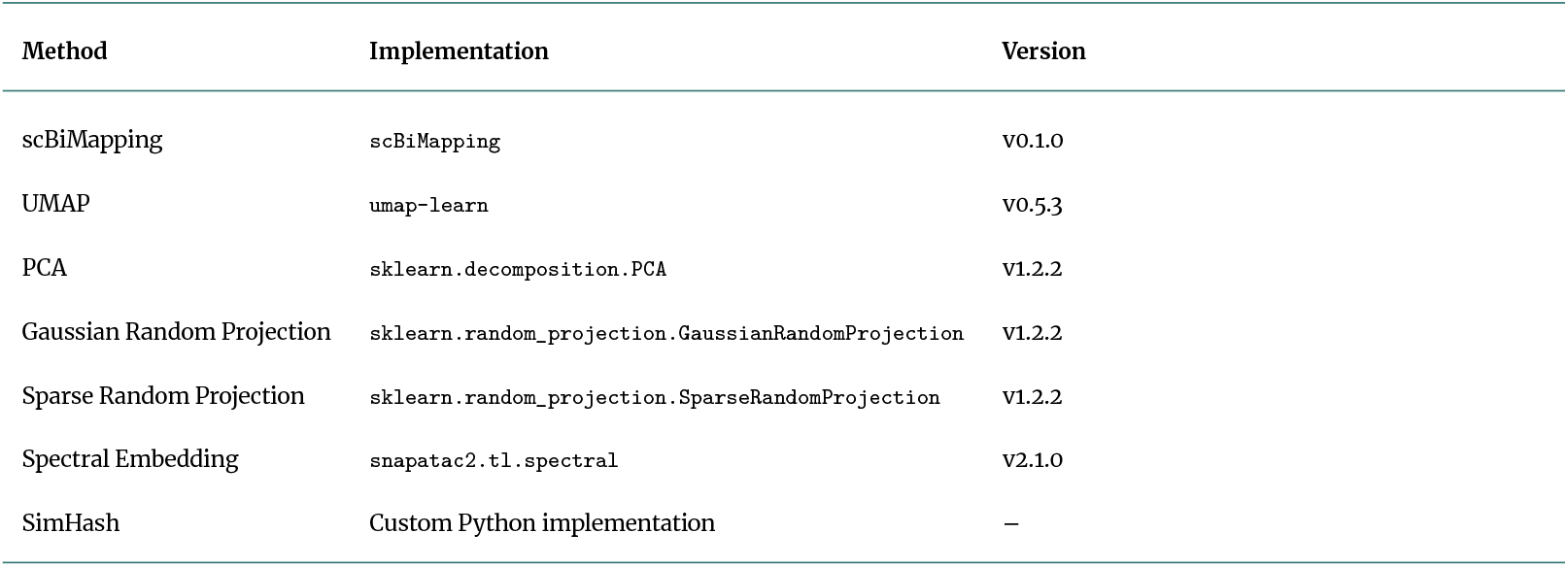
Summary of dimensionality reduction methods used in this study. Custom Python implementation of SimHash was used as no existing libraries suitable for analyzing biological sequences were found.

**Table S3.** Summary of k-nearest neighbor search methods used in this study.

| Method | Implementation | Version | Parameter |
| --- | --- | --- | --- |
| brute-force search | `sklearn.neighbors.NearestNeighbors` | v1.2.2 | `n_jobs = 64` |
| HNSW | `hnswlib` | v0.7.0 | `M = 512, ef_construction = 200, set_ef = 50, set_num_threads = 64` |
| PQ | `faiss` | v1.8.0 | `omp_set_num_threads = 64, index param = "PQ128x8"` |
| IVF-PQ | `faiss` | v1.8.0 | `omp_set_num_threads = 64, M = 128, nlist = 1024, nbits_per_idx = 8, nprobe = 300` |
| RPF | `rpforest` | v1.6 | `leaf_size = 50, no_trees = 100` |
| NNDescent | `pynndescent` | v0.5.12 | `n_jobs = 64, n_trees = 600, leaf_size = 200, index_n_neighbors = 50` |

**Table S4.**
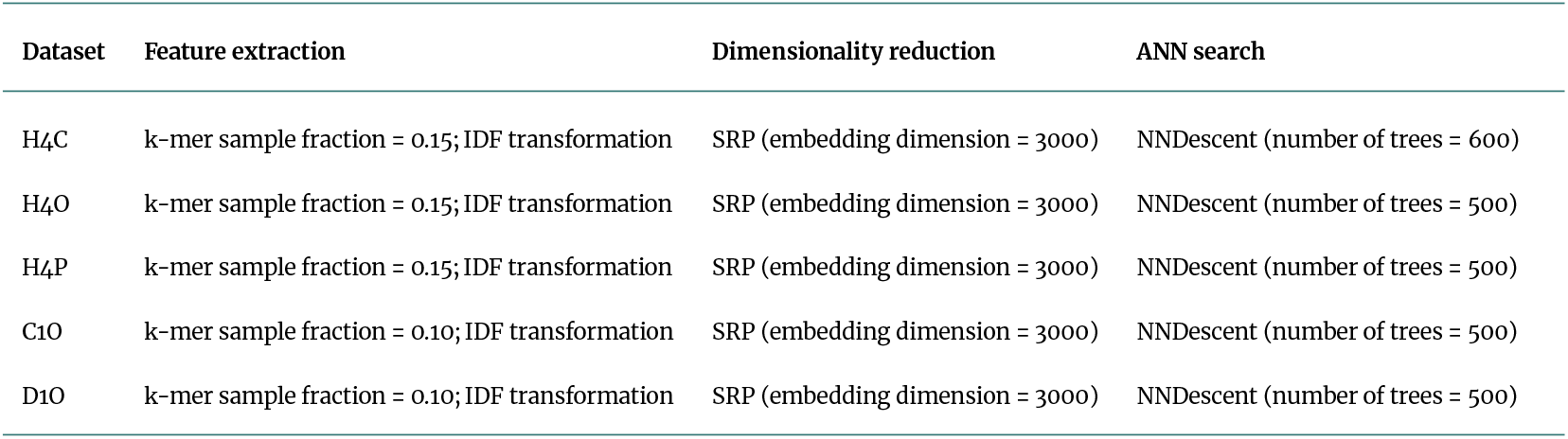
Fedrann parameters used for benchmarking.

## References

1. Sohn Ji, Nam JW. The present and future of de novo whole-genome assembly. Briefings in bioinformatics 2018;19(1):23–40.

2. Zhang JY, Zhang Y, Wang L, Guo F, Yun Q, Zeng T, et al. A single-molecule nanopore sequencing platform. bioRxiv 2024;p. 2024–08.

3. Nurk S, Koren S, Rhie A, Rautiainen M, Bzikadze AV, Mikheenko A, et al. The complete sequence of a human genome. Science 2022;376(6588):44–53.

4. Espinosa E, Bautista R, Larrosa R, Plata O. Advancements in long-read genome sequencing technologies and algorithms. Genomics 2024;p. 110842.

5. Koren S, Walenz BP, Berlin K, Miller JR, Bergman NH, Phillippy AM. Canu: scalable and accurate long-read assembly via adaptive k-mer weighting and repeat separation. Genome research 2017;27(5):722–736.

6. Shafin K, Pesout T, Lorig-Roach R, Haukness M, Olsen HE, Bosworth C, et al. Nanopore sequencing and the Shasta toolkit enable efficient de novo assembly of eleven human genomes. Nature biotechnology 2020;38(9):1044–1053.

7. Nie F, Ni P, Huang N, Zhang J, Wang Z, Xiao C, et al. De novo diploid genome assembly using long noisy reads. Nature Communications 2024;15(1):2964.

8. Rizzi R, Beretta S, Patterson M, Pirola Y, Previtali M, Della Vedova G, et al. Overlap graphs and de Bruijn graphs: data structures for de novo genome assembly in the big data era. Quantitative Biology 2019;7:278–292.

9. Altschul S. Gapped BLAST and PSI-BLAST: a new generation of protein database search programs. Bioinformatics 2012;298:3389.

10. Myers G. Efficient local alignment discovery amongst noisy long reads. In: International Workshop on Algorithms in Bioinformatics Springer; 2014. p. 52–67.

11. Li H. Minimap2: pairwise alignment for nucleotide sequences. Bioinformatics 2018;34(18):3094–3100.

12. Xiao CL, Chen Y, Xie SQ, Chen KN, Wang Y, Han Y, et al. MECAT: fast mapping, error correction, and de novo assembly for single-molecule sequencing reads. nature methods 2017;14(11):1072–1074.

13. Ruan J, Li H. Fast and accurate long-read assembly with wtdbg2. Nature methods 2020;17(2):155–158.

14. Kong T, Wang Y, Liu B. xRead: a coverage-guided approach for scalable construction of read overlapping graph. GigaScience 2025;14:giaf007.

15. Vinga S, Almeida J. Alignment-free sequence comparison—a review. Bioinformatics 2003;19(4):513–523.

16. Broder AZ. On the resemblance and containment of documents. In: Proceedings. Compression and Complexity of SEQUENCES 1997 (Cat. No. 97TB100171) IEEE; 1997. p. 21–29.

17. Berlin K, Koren S, Chin CS, Drake JP, Landolin JM, Phillippy AM. Assembling large genomes with single-molecule sequencing and locality-sensitive hashing. Nature biotechnology 2015;33(6):623–630.

18. Firtina C, Park J, Alser M, Kim JS, Cali DS, Shahroodi T, et al. BLEND: a fast, memory-efficient and accurate mechanism to find fuzzy seed matches in genome analysis. NAR Genomics and Bioinformatics 2023;5(1):lqad004.

19. Li H. Minimap and miniasm: fast mapping and de novo assembly for noisy long sequences. Bioinformatics 2016;32(14):2103–2110.

20. Greenberg GC. Analysis and applications of k-mer based methods in bioinformatics. PhD thesis, University of Illinois at Urbana-Champaign; 2023.

21. Charikar MS. Similarity estimation techniques from rounding algorithms. In: Proceedings of the thiry-fourth annual ACM symposium on Theory of computing; 2002. p. 380–388.

22. Ponsero AJ, Miller M, Hurwitz BL. Comparison of k-mer-based de novo comparative metagenomic tools and approaches. Microbiome research reports 2023;2(4):27.

23. Miller JR, Delcher AL, Koren S, Venter E, Walenz BP, Brownley A, et al. Aggressive assembly of pyrosequencing reads with mates. Bioinformatics 2008;24(24):2818–2824.

24. Zhang JY, Roberts H, Flores DS, Cutler AJ, Brown AC, Whalley JP, et al. Using de novo assembly to identify structural variation of eight complex immune system gene regions. PLoS computational biology 2021;17(8):e1009254.

25. Dunteman GH. Principal components analysis, vol. 69. Sage; 1989.

26. McInnes L, Healy J, Melville J. Umap: Uniform manifold approximation and projection for dimension reduction. arXiv preprint arXiv:180203426 2018;.

27. Dong W, Moses C, Li K. Efficient k-nearest neighbor graph construction for generic similarity measures. In: Proceedings of the 20th international conference on World wide web; 2011. p. 577–586.

28. Li J, Tian Y, Wang Y, Jin L. dna2bit: high performance genomic distance estimation software for microbial genome analysis. Frontiers in Microbiology 2024;15:1521181.

29. Malkov YA, Yashunin DA. Efficient and robust approximate nearest neighbor search using hierarchical navigable small world graphs. IEEE transactions on pattern analysis and machine intelligence 2018;42(4):824–836.

30. Jegou H, Douze M, Schmid C. Product quantization for nearest neighbor search. IEEE transactions on pattern analysis and machine intelligence 2010;33(1):117–128.

31. Yan D, Wang Y, Wang J, Wang H, Li Z. K-nearest neighbor search by random projection forests. IEEE Transactions on Big Data 2019;7(1):147–157.

32. Harris CR, Millman KJ, Van Der Walt SJ, Gommers R, Virtanen P, Cournapeau D, et al. Array programming with NumPy. Nature 2020;585(7825):357–362.

33. Virtanen P, Gommers R, Oliphant TE, Haberland M, Reddy T, Cournapeau D, et al. SciPy 1.0: fundamental algorithms for scientific computing in Python. Nature methods 2020;17(3):261–272.

34. Marçais G, Kingsford C. A fast, lock-free approach for efficient parallel counting of occurrences of k-mers. Bioinformatics 2011;27(6):764–770.

35. Ono Y, Hamada M, Asai K. PBSIM3: a simulator for all types of PacBio and ONT long reads. NAR genomics and bioinformatics 2022;4(4):lqac092.

36. Zook JM, Hansen NF, Olson ND, Chapman L, Mullikin JC, Xiao C, et al. A robust benchmark for detection of germline large deletions and insertions. Nature biotechnology 2020;38(11):1347–1355.

